# ATR is a master regulator of the oocyte meiotic prophase checkpoint

**DOI:** 10.64898/2026.10.13.758727

**Authors:** Jiwei Li, Xiaofei Jiao, Long Bai, Yuan Wang, Yimin Zhu, Yidan Liu, Lin-Yu Lu

## Abstract

During meiotic prophase, programmed double-strand breaks (DSBs) initiate meiotic recombination, and defects in this process result in persistent DSBs and chromosome asynapsis, triggering checkpoint-mediated oocyte elimination before primordial follicle formation. However, how the oocyte meiotic prophase checkpoint is regulated remains unclear. ATR is a DNA damage response kinase essential for male meiosis, but its role in female meiosis is poorly characterized. Here, we identify ATR as a master regulator of the oocyte meiotic prophase checkpoint. ATR loss fully rescues the survival of *Dmc1^-/-^*oocytes, which exhibit persistent DSBs and extensive chromosome asynapsis. ATR loss also fully rescues the survival of asynaptic *Spo11^-/-^* oocytes and *Trip13^-/-^* oocytes with aberrant checkpoint activation, demonstrating an essential role for ATR in the chromosome asynapsis checkpoint. Mechanistically, BRCA1 is required for ATR localization to unsynapsed chromosome axes, whereas ATR is dispensable for BRCA1 localization, establishing a BRCA1-ATR regulatory hierarchy at unsynapsed chromosome axes that is conserved between oocytes and spermatocytes. Together, these findings place ATR at the center of the oocyte meiotic prophase checkpoint and define a BRCA1-ATR signaling axis that links chromosome asynapsis to oocyte elimination.

## Introduction

Meiosis is a specialized cell division that generates haploid gametes for sexual reproduction. During meiotic prophase, programmed DNA double-strand breaks (DSBs) are generated and repaired by meiotic recombination using homologous chromosomes as templates. Concomitantly, homologous chromosomes undergo synapsis through formation of the synaptonemal complex(1). Meiotic recombination generates crossovers between homologous chromosomes, which are essential for their proper segregation during the first meiotic division(2). Defects in meiotic recombination can result in persistent DSBs and chromosome asynapsis, compromising genome integrity and gamete quality(3). To prevent the production of defective gametes, meiotic recombination is closely monitored by meiotic prophase checkpoints that eliminate cells with recombination defects(4–6).

In females, recombination-defective oocytes are eliminated before the establishment of the primordial follicle pool(7–11), but the mechanisms underlying the oocyte meiotic prophase checkpoint remain incompletely understood. The CHK2-dependent DNA damage checkpoint is well established in the elimination of oocytes exposed to exogenous DNA damage, and CHK2 is also required for the efficient elimination of recombination-defective oocytes(8,9). However, chromosome asynapsis, which often coexists with persistent DSBs in recombination-defective oocytes, can trigger oocyte elimination independently of persistent DSBs(12). We recently identified BRCA1, a canonical DNA damage response protein in somatic cells, as a critical regulator of the oocyte chromosome asynapsis checkpoint(13). Unlike in somatic cells, BRCA1 does not specifically localize to DSBs during meiotic prophase but instead localizes to unsynapsed chromosome axes together with HORMAD1 and HORMAD2, which monitor chromosome synapsis(13–17). *Brca1^-/-^*partially rescues the survival of recombination-defective *Dmc1^-/-^* oocytes and asynaptic *Spo11^-/-^* oocytes, supporting an important role for the BRCA1-dependent chromosome asynapsis checkpoint in eliminating recombination-defective oocytes(13).

Aberrant activation of the BRCA1-dependent chromosome asynapsis checkpoint also leads to oocyte elimination(18). TRIP13 is an AAA+ ATPase required for the removal of HORMAD1 and HORMAD2 from synapsed chromosome axes during meiotic prophase(19–22). In *Trip13^-/-^* oocytes, DSB repair is delayed but completed before oocyte elimination, whereas HORMAD1 and HORMAD2 are aberrantly retained on synapsed chromosome axes, leading to persistent BRCA1 localization on synapsed chromosome axes(18). *Trip13^-/-^*oocytes are eliminated before the establishment of the primordial follicle pool(19,20). Importantly, *Brca1^-/-^* partially rescues the survival of *Trip13^-/-^*oocytes, further suggesting that aberrant activation of the chromosome asynapsis checkpoint, rather than persistent DNA damage, is responsible for oocyte elimination in *Trip13^-/-^* females(18). Therefore, the chromosome asynapsis checkpoint must be precisely regulated to prevent aberrant activation and inappropriate oocyte elimination.

How BRCA1 regulates the oocyte chromosome asynapsis checkpoint remains unclear, but ATR is a likely downstream mediator(13,18). ATR is a DNA damage response kinase essential for male meiosis(23–25). ATR localizes to the unsynapsed axes of male sex chromosomes and promotes meiotic sex chromosome inactivation(23,24). ATR also regulates meiotic recombination in males(24,25), although the precise mechanism remains unclear. In contrast, the role of ATR in female meiosis has been far less studied, largely due to the difficulty in effectively depleting ATR during female meiotic prophase. Meiotic recombination is largely normal in *Atr^s/s^* mutant mice carrying a mutation identified in patients with Seckel syndrome, but the hypomorphic nature of this mutation precludes concluding that ATR is dispensable for female meiotic recombination(26). Importantly, the role of ATR in the oocyte meiotic prophase checkpoint remains largely unknown.

In this study, we identify ATR as a master regulator of the oocyte meiotic prophase checkpoint and a critical downstream mediator of BRCA1 in the chromosome asynapsis checkpoint. ATR is essential for checkpoint activation in response to unsynapsed chromosomes as well as for aberrant checkpoint activation on synapsed chromosomes, but is dispensable for oocyte elimination in response to ionizing radiation (IR)-induced DNA damage. We further demonstrate that BRCA1 regulates ATR localization to unsynapsed chromosome axes, establishing a BRCA1-ATR regulatory hierarchy conserved between oocytes and spermatocytes.

## Results

### ATR localizes to unsynapsed chromosome axes in a BRCA1-dependent manner during oocyte meiotic prophase

To investigate the role of ATR in female meiosis, we first examined its localization during meiotic prophase in oocytes. In our previous study, ATR signals were reliably detected in oocytes at embryonic day (E) 16.5 but not at postnatal day (PD) 0(13), leading us to speculate that ATR might transiently localize to unsynapsed chromosome axes during meiotic prophase. Using a new lot of ATR antibody that produced more robust signals, we reliably detected ATR in both embryonic and postnatal oocytes, allowing us to examine its localization throughout meiotic prophase. ATR was detected on unsynapsed chromosome axes at leptotene, disappeared from SYCP1-positive regions as homologous chromosomes underwent synapsis at zygotene, and was absent from fully synapsed chromosome axes at pachytene (Figure 1A). At all examined stages of meiotic prophase, ATR colocalized with HORMAD1 (Figure 1B), consistent with its specific localization to unsynapsed chromosome axes.

**Figure 1.**
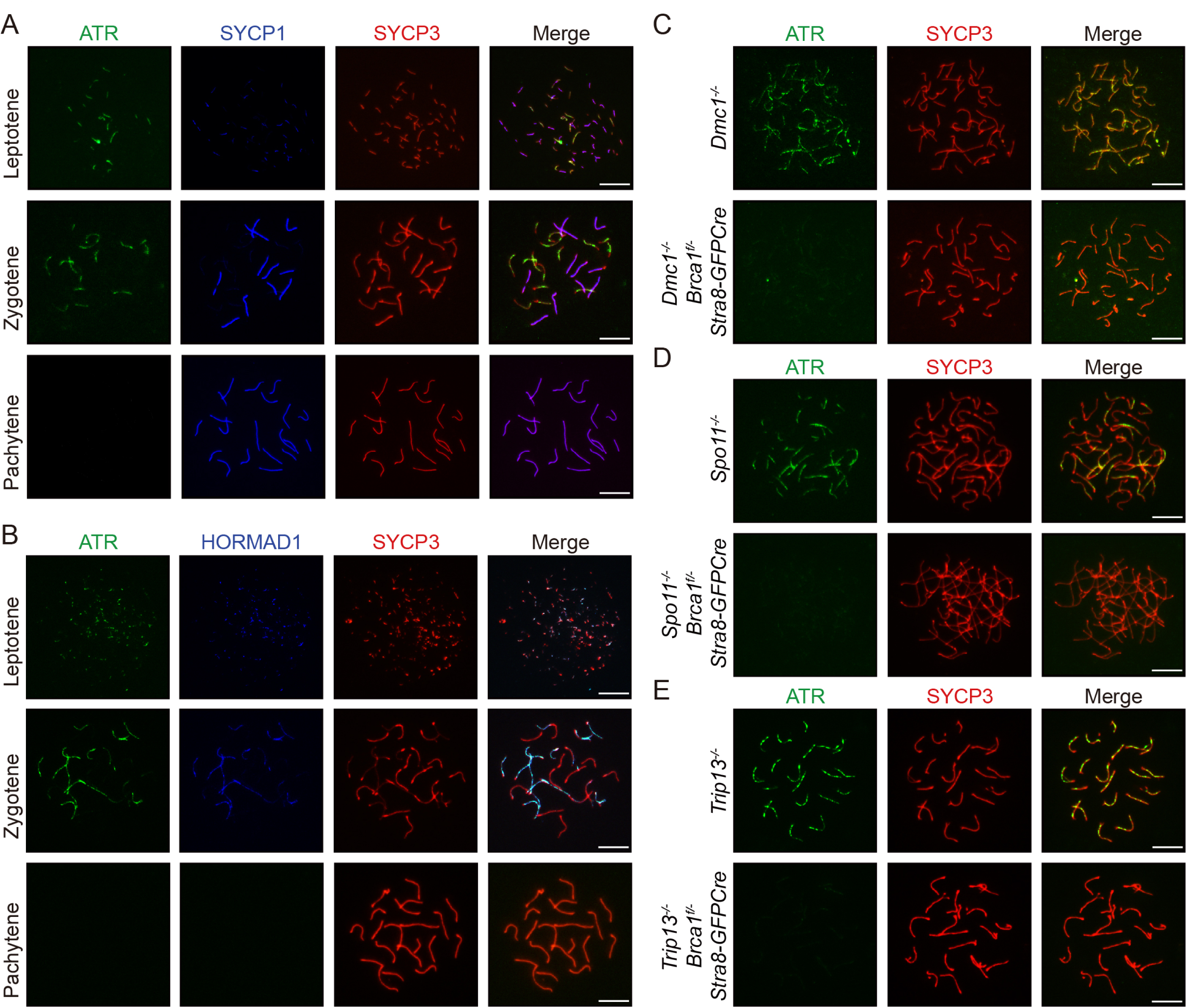
ATR localizes to unsynapsed chromosome axes in a BRCA1-dependent manner during oocyte meiotic prophase. (A, B) Representative images of ATR, SYCP1, and SYCP3 (A) and ATR, HORMAD1, and SYCP3 (B) staining in chromosome spreads of oocytes at leptotene, zygotene, and pachytene from WT females. Scale bars, 10 μm. (C, D) Representative images of ATR and SYCP3 staining in chromosome spreads of *Dmc1^-/-^* and *Dmc1^-/-^ Brca1^f/-^ Stra8-GFPCre* oocytes (C), *Spo11^-/-^* and *Spo11^-/-^ Brca1^f/-^ Stra8-GFPCre* oocytes (D) at zygotene stages. Scale bars, 10 μm. (E) Representative images of ATR and SYCP3 staining in chromosome spreads of *Trip13^-/-^* and *Trip13^-/-^ Brca1^f/-^ Stra8-GFPCre* oocytes at pachytene stages. Scale bars, 10 μm.

We next examined ATR localization in several mutant oocytes in which the oocyte meiotic prophase checkpoint is activated. In recombination-defective *Dmc1^-/-^*oocytes, chromosome synapsis failed globally, and ATR localized to the unsynapsed axes of all chromosomes (Figure 1C). In asynaptic *Spo11^-/-^* oocytes, in which programmed DSBs are not generated, ATR also localized to unsynapsed chromosome axes, indicating that this localization is independent of programmed DSBs (Figure 1D). In *Trip13^-/-^* oocytes, in which chromosome synapsis is normal but the chromosome asynapsis checkpoint is aberrantly activated, ATR was retained on the axes of fully synapsed chromosomes (Figure 1E). Therefore, ATR localization to chromosome axes closely correlates with activation of the oocyte chromosome asynapsis checkpoint.

Our previous study showed that BRCA1, which also localizes to chromosome axes in a DSB-independent manner, is a key regulator of the chromosome asynapsis checkpoint(13,18). *Brca1^-/-^* partially rescues the survival of *Dmc1^-/-^*, *Spo11^-/-^*, and *Trip13^-/-^* oocytes. Our previous study also suggested that ATR is an important downstream mediator of BRCA1 in the chromosome asynapsis checkpoint(13,18). To further test this possibility, we used *Brca1^f/-^ Stra8-GFPCre* mice, in which BRCA1 is effectively depleted before chromosome asynapsis checkpoint activation(13), and examined ATR localization in these oocytes. *Brca1^-/-^* dramatically disrupted ATR localization to chromosome axes in *Dmc1^-/-^*, *Spo11^-/-^*, and *Trip13^-/-^* oocytes (Figure 1C-E). Therefore, BRCA1 is required for ATR localization to chromosome axes, suggesting that ATR functions downstream of BRCA1 in the oocyte chromosome asynapsis checkpoint.

### ATR is required for female fertility

To investigate the role of ATR in female meiosis, we generated *Atr^f/-^ Stra8-GFPCre* mice, in which ATR was effectively depleted by PD0 (Figure S1A-B). ATR depletion did not alter BRCA1 localization to unsynapsed chromosome axes (Figure S1C), further suggesting that BRCA1 acts upstream of ATR at these axes. Previous studies have shown that ATR is required for male meiotic recombination(24,25). However, no obvious defects in meiotic prophase progression, chromosome synapsis, or crossover formation, as assessed by MLH1 foci, were observed in *Atr^f/-^ Stra8-GFPCre* females (Figure S1D-G), indicating largely normal meiotic recombination.

Despite largely normal meiotic recombination and morphologically normal ovaries, *Atr^f/-^ Stra8-GFPCre* females were infertile (Figure 2A-B). During the first meiotic division, *Atr^-/-^* oocytes formed normal meiotic spindles with chromosomes properly aligned at the metaphase plate (Figure 2C-D). Polar body extrusion rates were also comparable between control and *Atr^-/-^* oocytes (Figure 2E-F). After fertilization of *Atr^-/-^* oocytes with wild-type sperm, the resulting zygotes developed to the two-cell stage without apparent delay at E1.5 (Figure 2G-H). However, subsequent development was severely impaired, and no blastocysts were observed at E3.5 (Figure 2G-H). Therefore, *Atr^-/-^* oocytes failed to support normal preimplantation embryonic development, accounting for the infertility of *Atr^f/-^ Stra8-GFPCre* females.

**Figure 2.**
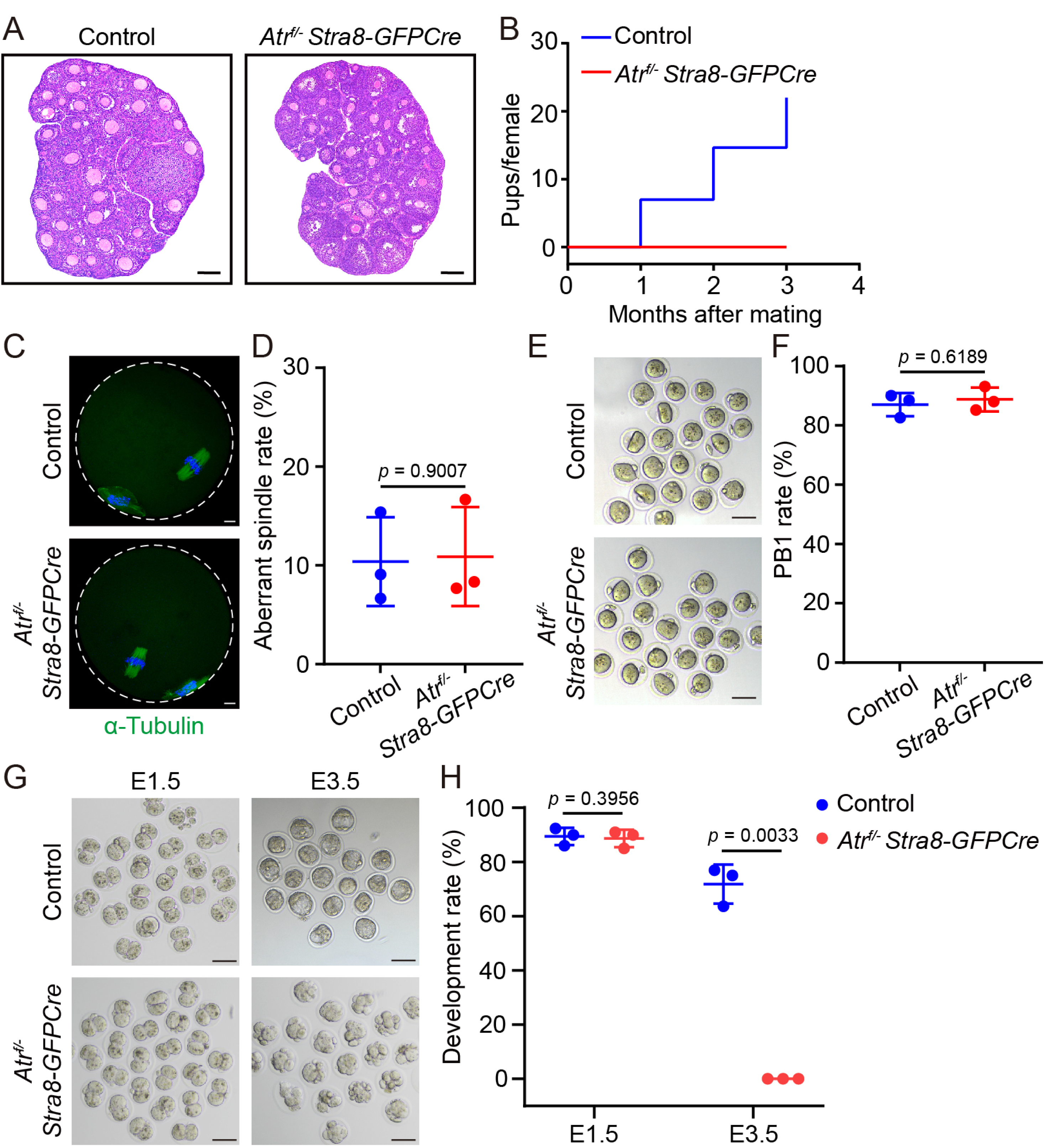
ATR is required for female fertility. (A) Representative images of H&E staining of ovarian sections obtained from control and *Atr^f/-^ Stra8-GFPCre* mice at PD21. Scale bars, 100 μm. (B) Cumulative number of pups per female obtained by mating control and *Atr^f/-^ Stra8-GFPCre* adult female mice with WT males for 3 consecutive months. (C) Representative images of spindle morphology and chromosome alignment in oocytes from control and *Atr^f/-^ Stra8-GFPCre* females. Scale bars, 10 μm. (D) Aberrant spindle rate of in vitro matured oocytes from control and *Atr^f/-^ Stra8-GFPCre* females. (E) Representative images of control and *Atr^f/-^ Stra8-GFPCre* oocytes after first polar body (PB1) extrusion. Scale bars, 100 μm. (F) Statistics of PB1 extrusion rate of control and *Atr^f/-^ Stra8-GFPCre* oocytes. (G) Representative images of embryos collected from control and *Atr^f/-^ Stra8- GFPCre* females at E1.5/E3.5 after mating with WT males. Scale bars, 100 μm. (H) Development rate of embryos collected from control and *Atr^f/-^ Stra8-GFPCre* females at E1.5/E3.5 after mating with WT males. All statistical data are presented as means ± SD. *P* values were determined by two-tailed unpaired Student’s t tests.

### ATR is a master regulator of the oocyte meiotic prophase checkpoint

We next investigated the role of ATR in the oocyte meiotic prophase checkpoint. To this end, we generated *Dmc1^-/-^ Atr^f/-^ Stra8-GFPCre* mice and examined whether *Atr^-/-^* rescues the survival of recombination-defective *Dmc1^-/-^* oocytes, as previously observed with *Brca1^-/-^*(13). Due to oocyte elimination before the establishment of the primordial follicle pool, no follicles were present in the atrophied ovaries of *Dmc1^-/-^* females at PD21 (Figure 3A). In contrast, the ovaries of *Dmc1^-/-^ Atr^f/-^ Stra8-GFPCre* females were morphologically similar to those of controls at PD21 (Figure 3A). Total and primordial follicle numbers in *Dmc1^-/-^ Atr^f/-^ Stra8-GFPCre* ovaries were not significantly different from those in control ovaries at PD21 (Figure 3B). Total follicle numbers in adult *Dmc1^-/-^ Atr^f/-^ Stra8-GFPCre* ovaries remained comparable to those in control ovaries at PD60 (Figure 3C-D). Together, these results indicate that *Atr^-/-^* fully rescues the survival of *Dmc1^-/-^* oocytes. In our previous study, *Brca1^-/-^* only partially rescued the survival of *Dmc1^-/-^*oocytes(13). Consistent with this difference, *Dmc1^-/-^ Atr^f/-^ Stra8-GFPCre* ovaries contained substantially more total and primordial follicles than *Dmc1^-/-^ Brca1^f/-^ Stra8-GFPCre* ovaries at PD21 and PD60 (Figure 3B, D). Therefore, *Atr^-/-^* is more effective than *Brca1^-/-^*in rescuing the survival of *Dmc1^-/-^* oocytes, supporting a central role for ATR in the oocyte meiotic prophase checkpoint.

**Figure 3.**
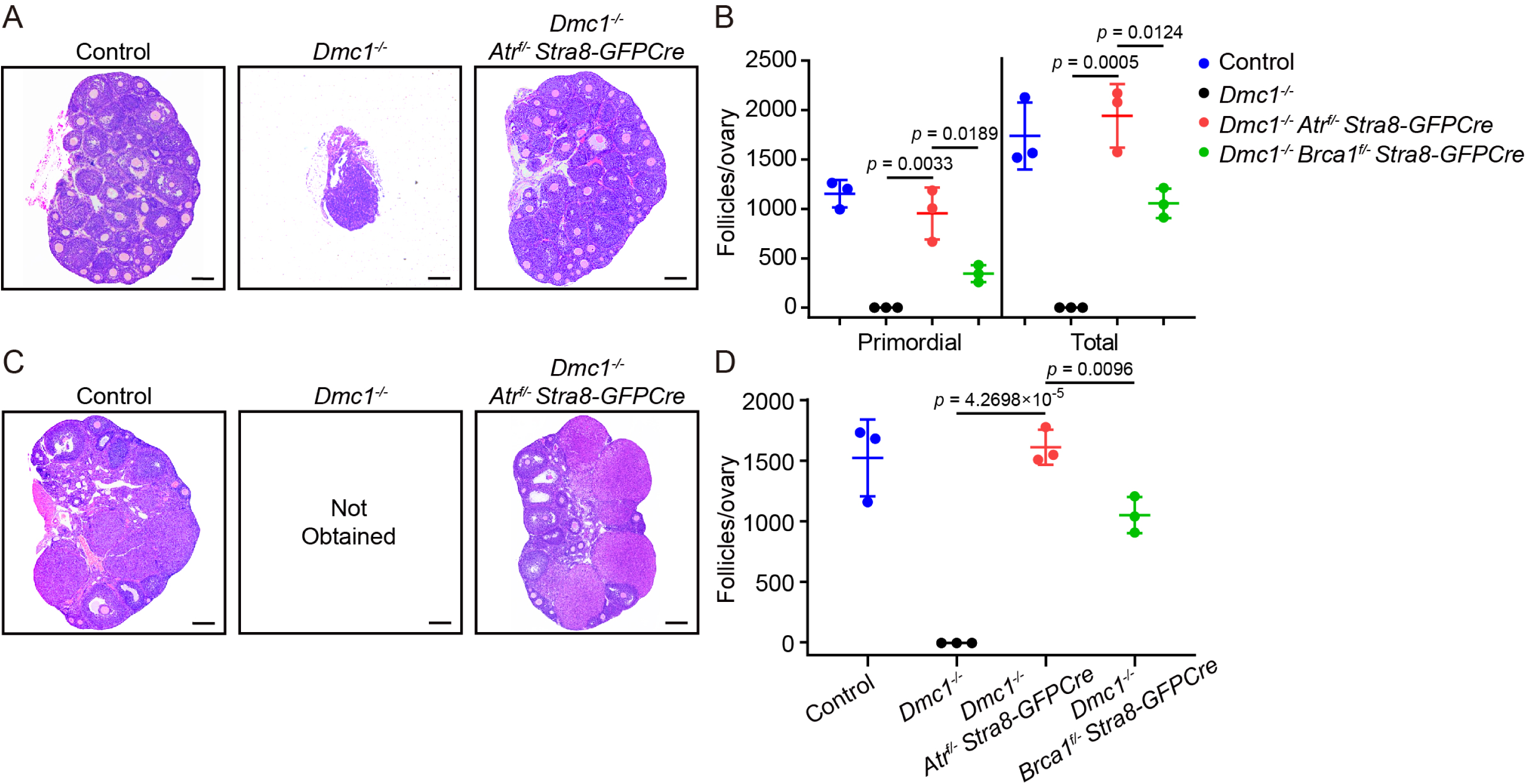
ATR is a master regulator of the oocyte meiotic prophase checkpoint. (A) Representative images of H&E staining of ovarian sections from control, *Dmc1^-/-^*, and *Dmc1^-/-^ Atr^f/-^ Stra8-GFPCre* mice at PD21. Scale bars, 100 μm. (B) Quantification of total and primordial follicles in control, *Dmc1^-/-^*, *Dmc1^-/-^ Atr^f/-^ Stra8-GFPCre*, and *Dmc1^-/-^ Brca1^f/-^ Stra8-GFPCre* mice at PD21. (C) Representative images of H&E staining of ovarian sections from control, *Dmc1^-/-^*, and *Dmc1^-/-^ Atr^f/-^ Stra8-GFPCre* mice at PD60. Scale bars, 100 μm. (D) Quantification of total follicles in control, *Dmc1^-/-^*, *Dmc1^-/-^ Atr^f/-^ Stra8-GFPCre*, and *Dmc1^-/-^ Brca1^f/-^ Stra8-GFPCre* mice at PD60. All statistical data are presented as means ± SD. *P* values were determined by two-tailed unpaired Student’s t tests. The *Dmc1^-/-^ Brca1^f/-^ Stra8-GFPCre* mice data were replotted from our previous study for comparison(13).

### ATR is essential for the BRCA1-dependent chromosome asynapsis checkpoint

Our findings in *Dmc1^-/-^* oocytes establish an essential role for ATR in the oocyte meiotic prophase checkpoint, but these oocytes exhibit both persistent DSBs and extensive chromosome asynapsis. To directly examine the role of ATR in the chromosome asynapsis checkpoint, we generated *Spo11^-/-^ Atr^f/-^ Stra8-GFPCre* mice and examined whether *Atr^-/-^* rescues the survival of asynaptic *Spo11^-/-^*oocytes, which lack programmed DSBs. The ovaries of *Spo11^-/-^ Atr^f/-^ Stra8-GFPCre* females were morphologically similar to those of controls at PD60 (Figure 4A). Total and primordial follicle numbers in *Spo11^-/-^ Atr^f/-^ Stra8-GFPCre* ovaries were comparable to those in control ovaries and substantially higher than those in *Spo11^-/-^ Brca1^f/-^ Stra8-GFPCre* ovaries analyzed in our previous study (Figure 4B)(13). Therefore, *Atr^-/-^*fully rescues the survival of asynaptic *Spo11^-/-^* oocytes, demonstrating that ATR is essential for the chromosome asynapsis checkpoint.

**Figure 4.**
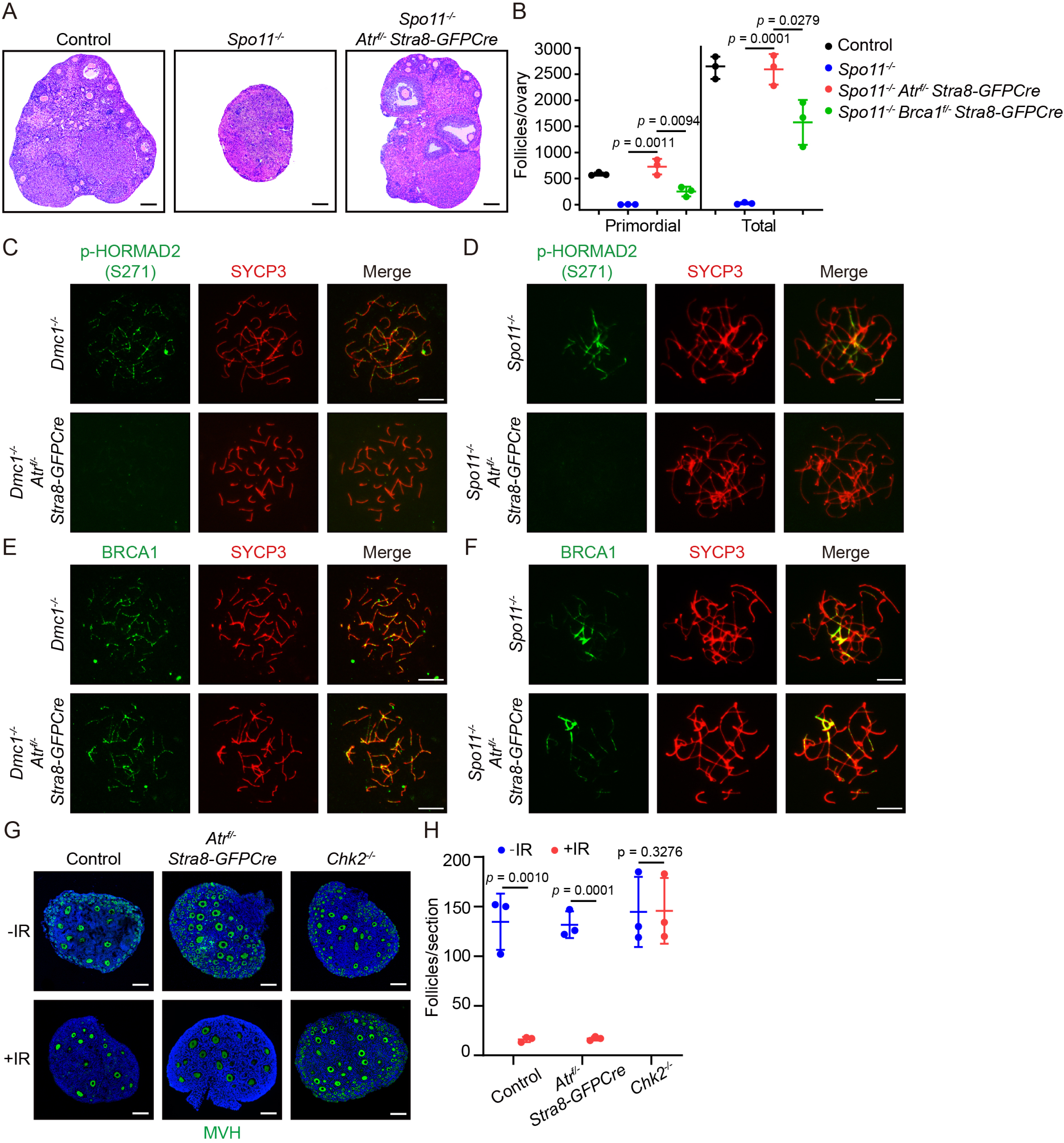
ATR is essential for the BRCA1-dependent chromosome asynapsis checkpoint. (A) Representative images of H&E staining of ovarian sections from control, *Spo11^-/-^*, and *Spo11^-/-^ Atr^f/-^ Stra8-GFPCre* mice at PD60. Scale bars, 100 μm. (B) Quantification of total and primordial follicles in control, *Spo11^-/-^*, *Spo11^-/-^ Atr^f/-^ Stra8-GFPCre*, and *Spo11^-/-^ Brca1^f/-^ Stra8-GFPCre* mice at PD60. (C, D) Representative images of p-HORMAD2 (S271) and SYCP3 staining in chromosome spreads of *Dmc1^-/-^*, *Dmc1^-/-^ Atr^f/-^ Stra8-GFPCre* (C), *Spo11^-/-^*, and *Spo11^-/-^ Atr^f/-^ Stra8-GFPCre* (D) oocytes at zygotene stages. Scale bars, 10 μm. (E, F) Representative images of BRCA1 and SYCP3 staining in chromosome spreads of *Dmc1^-/-^*, *Dmc1^-/-^ Atr^f/-^ Stra8-GFPCre* (E), *Spo11^-/-^*, and *Spo11^-/-^ Atr^f/-^ Stra8-GFPCre* (F) oocytes at zygotene stages. Scale bars, 10 μm. (G) Representative images of MVH staining in ovarian sections at PD9 from control, *Atr^f/-^ Stra8-GFPCre*, and *Chk2^-/-^* females with or without 0.45 Gy IR exposure at PD2. Scale bars, 10 μm. (H) Quantification of MVH-positive oocytes per section at PD9 from control, *Atr^f/-^ Stra8-GFPCre*, and *Chk2^-/-^* females with or without 0.45 Gy IR exposure at PD2. All statistical data are presented as means ± SD. *P* values were determined by two-tailed unpaired Student’s t tests. The *Spo11^-/-^ Brca1^f/-^ Stra8-GFPCre* mice data were replotted from our previous study for comparison(13).

ATR loss markedly reduced phosphorylation of HORMAD2, a known ATR substrate(23), at unsynapsed chromosome axes in both *Dmc1^-/-^* oocytes and *Spo11^-/-^*oocytes (Figure 4C-D). However, ATR loss did not affect BRCA1 localization to unsynapsed chromosome axes in either *Dmc1^-/-^* or *Spo11^-/-^* oocytes (Figure 4E-F). Together with our finding that *Brca1^-/-^* dramatically disrupted ATR localization to unsynapsed chromosome axes in both mutants, these results further suggest that ATR acts downstream of BRCA1 in the oocyte chromosome asynapsis checkpoint.

To determine whether ATR participates in the oocyte DNA damage checkpoint, we examined whether ATR loss renders oocytes resistant to IR-induced elimination. To this end, we exposed *Atr^f/-^ Stra8-GFPCre* females to 0.45 Gy IR, which effectively eliminated the majority of follicles in control females(8,9). Loss of CHK2 rendered oocytes resistant to 0.45 Gy IR, with no significant reduction in follicle numbers (Figure 4G-H). In contrast, 0.45 Gy IR effectively eliminated the majority of follicles in *Atr^f/-^ Stra8-GFPCre* females, and the numbers of remaining follicles were comparable to those in irradiated controls (Figure 4G-H). Therefore, unlike CHK2, ATR is dispensable for oocyte elimination in response to IR-induced DNA damage. However, these results do not exclude the possibility that ATR participates in the oocyte DNA damage checkpoint in response to persistent programmed DSBs.

### Aberrant ATR retention on synapsed chromosome axes triggers chromosome asynapsis checkpoint activation

We previously showed that retention of BRCA1 on synapsed chromosome axes in *Trip13^-/-^* oocytes leads to aberrant activation of the chromosome asynapsis checkpoint and oocyte elimination(18). Because ATR was also retained on synapsed chromosome axes in a BRCA1-dependent manner, we next examined whether ATR is required for aberrant checkpoint activation in *Trip13^-/-^* oocytes. To this end, we generated *Trip13^-/-^ Atr^f/-^ Stra8-GFPCre* mice and examined whether *Atr^-/-^* rescues the survival of *Trip13^-/-^* oocytes. In contrast to the atrophied ovaries of *Trip13^-/-^* females, the ovaries of *Trip13^-/-^ Atr^f/-^ Stra8-GFPCre* females were morphologically normal at PD21 (Figure 5A). Total and primordial follicle numbers in *Trip13^-/-^ Atr^f/-^ Stra8-GFPCre* ovaries were comparable to those in control ovaries and substantially higher than those in *Trip13^-/-^ Brca1^f/-^ Stra8-GFPCre* ovaries analyzed in our previous study (Figure 5B)(18). Therefore, *Atr^-/-^* fully rescues the survival of *Trip13^-/-^* oocytes, demonstrating that ATR is required for aberrant chromosome asynapsis checkpoint activation. Consistent with these findings, ATR loss markedly reduced HORMAD2 phosphorylation but did not affect BRCA1 localization to synapsed chromosome axes in *Trip13^-/-^* oocytes (Figure 5C-D), further suggesting that ATR functions downstream of BRCA1 in the oocyte chromosome asynapsis checkpoint.

**Figure 5.**
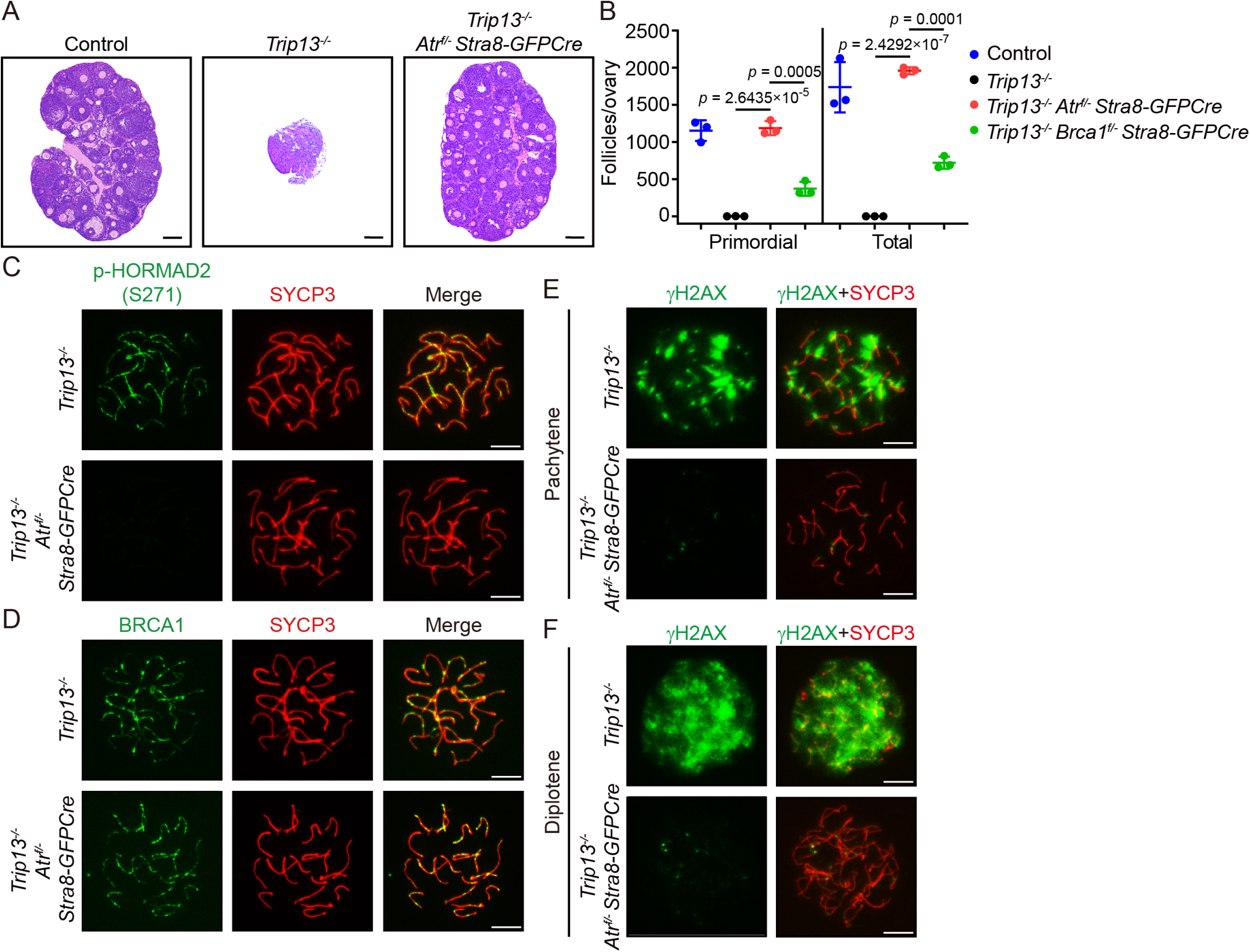
Aberrant ATR retention on synapsed chromosome axes triggers chromosome asynapsis checkpoint activation. (A) Representative images of H&E staining of ovarian sections from control, *Trip13^-/-^*, and *Trip13^-/-^ Atr^f/-^ Stra8-GFPCre* mice at PD21. Scale bars, 100 μm. (B) Quantification of total and primordial follicles in control, *Trip13^-/-^*, *Trip13^-/-^ Atr^f/-^ Stra8-GFPCre*, and *Trip13^-/-^ Brca1^f/-^ Stra8-GFPCre* mice at PD21. (C, D) Representative images of p-HORMAD2 (S271) and SYCP3 staining (C) and BRCA1 and SYCP3 staining (D) in chromosome spreads of *Trip13^-/-^* and *Trip13^-/-^ Atr^f/-^ Stra8-GFPCre* oocytes at pachytene stages. Scale bars, 10 μm. (E, F) Representative images of γH2AX and SYCP3 staining in chromosome spreads of *Trip13^-/-^*and *Trip13^-/-^ Atr^f/-^ Stra8-GFPCre* oocytes at pachytene (E) and diplotene (F) stages. Scale bars, 10 μm. All statistical data are presented as means ± SD. *P* values were determined by two-tailed unpaired Student’s t tests. The *Trip13^-/-^ Brca1^f/-^ Stra8-GFPCre* mice data were replotted from our previous study for comparison(18).

Based on the strong γH2AX signals observed in *Trip13^-/-^*oocytes before their elimination (Figure 5E-F), it was previously thought that persistent DSBs triggered their elimination. However, we recently showed that DSB repair is completed before *Trip13^-/-^* oocytes are eliminated, arguing against persistent DNA damage as the trigger, although the origin of the strong γH2AX signals remained unclear(18). In sharp contrast, very little γH2AX signal was detected in *Trip13^-/-^ Atr^f/-^ Stra8-GFPCre* oocytes (Figure 5E-F). Given the established role of ATR in H2AX phosphorylation(23–25), these results suggest that aberrantly retained ATR is responsible for the strong γH2AX signals in *Trip13^-/-^* oocytes. Therefore, the strong γH2AX signals in *Trip13^-/-^* oocytes likely reflect aberrant ATR-dependent checkpoint signaling rather than persistent DSBs, further supporting the conclusion that these oocytes are eliminated through aberrant activation of the chromosome asynapsis checkpoint.

### BRCA1 regulates ATR localization to unsynapsed chromosome axes in spermatocytes

In spermatocytes, both BRCA1 and ATR localize to unsynapsed chromosome axes in the XY body(23,24,27,28). Previous studies have shown that BRCA1 deficiency disrupts ATR localization to unsynapsed X and Y chromosome axes and that ATR deficiency also disrupts BRCA1 localization(23,27,28), suggesting reciprocal regulation between BRCA1 and ATR. This differs from our findings in oocytes, in which ATR loss does not affect BRCA1 localization to unsynapsed chromosome axes (Figure 4E-F). To further clarify the regulatory relationship between BRCA1 and ATR in spermatocytes, we used *Atr^f/-^ Stra8-GFPCre* males, in which ATR depletion begins in differentiating spermatogonia. *Atr^f/-^ Stra8-GFPCre* males were infertile, with no sperm detected in the epididymides (Figure S2A). Their testes were significantly smaller than those of control males, and no haploid spermatids were observed in testis sections (Figure S2B-D). ATR signals were undetectable at all examined stages of meiotic prophase in *Atr^f/-^ Stra8-GFPCre* males (Figure S2E-F), demonstrating efficient ATR depletion in spermatocytes. Meiotic progression was arrested at mid-pachytene (Figure S2G-H), consistent with the established roles of ATR in meiotic recombination and meiotic sex chromosome inactivation during male meiotic prophase.

As in oocytes, both BRCA1 and ATR localized to unsynapsed chromosome axes in zygotene spermatocytes, and their localization was not restricted to the X and Y chromosomes (Figure 6A-B). Consistent with our findings in oocytes, BRCA1 localization to unsynapsed chromosome axes was unaffected in zygotene spermatocytes from *Atr^f/-^ Stra8-GFPCre* males (Figure 6A). To determine whether BRCA1 regulates ATR localization to unsynapsed chromosome axes, we examined *Brca1^-/-^ 53bp1^-/-^* males, in which loss of 53BP1 rescues the embryonic lethality of *Brca1^-/-^*mice(29). ATR signals were absent from unsynapsed chromosome axes in zygotene spermatocytes from *Brca1^-/-^ 53bp1^-/-^* males (Figure 6B), indicating that BRCA1 is required for ATR localization to these axes. Together, these observations suggest that ATR functions downstream of BRCA1 at unsynapsed chromosome axes in spermatocytes. Both BRCA1 and ATR promote RAD51 and DMC1 loading during meiotic recombination (Figure 6C-F). Interestingly, *Atr^f/-^ Stra8-GFPCre* males showed a milder reduction in RAD51 and DMC1 foci than that observed in *Brca1^-/-^ 53bp1^-/-^* males (Figure 6C-F). Therefore, although BRCA1 regulates ATR localization to unsynapsed chromosome axes in spermatocytes, these results suggest that BRCA1 promotes recombinase loading partly independently of ATR.

**Figure 6.**
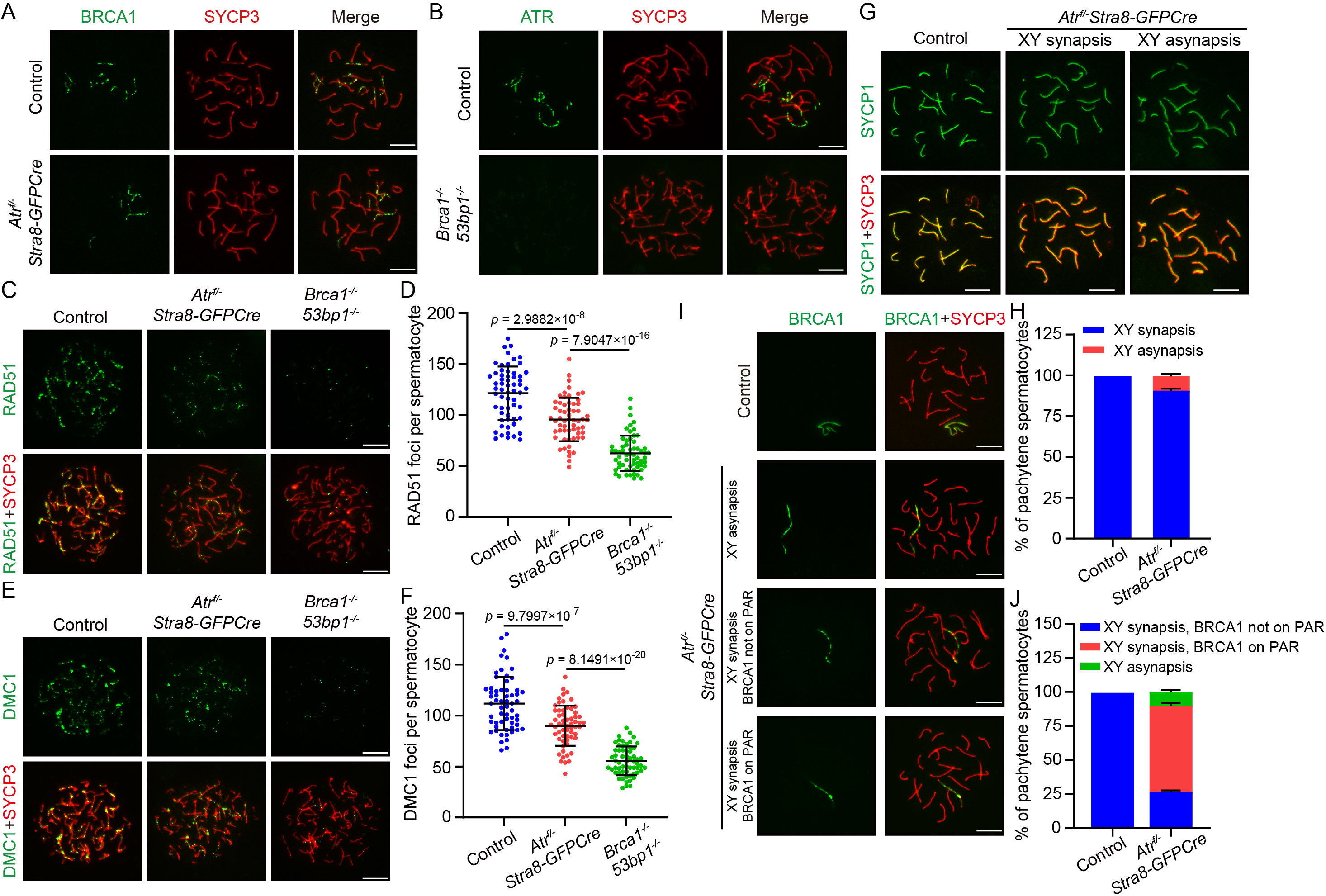
BRCA1 regulates ATR localization to unsynapsed chromosome axes in spermatocytes. (A) Representative images of BRCA1 and SYCP3 staining in chromosome spreads of control and *Atr^f/-^ Stra8-GFPCre* spermatocytes at zygotene stages. Scale bars, 10 μm. (B) Representative images of ATR and SYCP3 staining in chromosome spreads of control and *Brca1^-/-^ 53bp1^-/-^*spermatocytes at zygotene stages. Scale bars, 10 μm. (C, E) Representative images of RAD51 and SYCP3 staining (C) and DMC1 and SYCP3 staining (E) of control, *Atr^f/-^ Stra8-GFPCre*, and *Brca1^−/−^ 53bp1^−/−^* spermatocytes at zygotene stages. Scale bars, 10 μm. (D, F) Quantification of RAD51 foci (D) and DMC1 foci (F) in chromosome spreads of control, *Atr^f/-^ Stra8-GFPCre*, and *Brca1^−/−^ 53bp1^−/−^* spermatocytes at zygotene stages. (G) Representative images of SYCP1 and SYCP3 staining in chromosome spreads of control and *Atr^f/-^ Stra8-GFPCre* spermatocytes at pachytene stages. Scale bars, 10 μm. (H) Proportions of two spermatocyte classes with synapsed or asynapsed XY chromosomes at the pachytene stage in control and *Atr^f/-^ Stra8-GFPCre* mice. (I) Representative images of BRCA1 and SYCP3 staining in chromosome spreads of control and *Atr^f/-^ Stra8-GFPCre* spermatocytes at pachytene stages. Scale bars, 10 μm. (J) Proportions of three classes of spermatocytes, categorized by XY-chromosome synapsis status and BRCA1 localization at the pachytene stage in control and *Atr^f/-^ Stra8-GFPCre* mice. All statistical data are presented as means ± SD. *P* values were determined by two-tailed unpaired Student’s t tests.

Previous studies have reported that ATR loss disrupts BRCA1 localization to unsynapsed X and Y chromosome axes(23), which appears inconsistent with our findings in zygotene spermatocytes. We therefore examined BRCA1 localization to unsynapsed X and Y chromosome axes in *Atr^f/-^ Stra8-GFPCre* males. Because meiotic progression was arrested at mid-pachytene, no fully developed XY body was observed (Figure S2G). However, synapsis between the X and Y chromosomes at the pseudoautosomal region (PAR) was observed in most mid-pachytene spermatocytes (Figure 6G-H). Consistent with previous studies(23), ATR loss did not affect HORMAD1 or HORMAD2 localization to unsynapsed X and Y chromosome axes but dramatically reduced HORMAD2 phosphorylation and γH2AX signals (Figure S3A-D). In contrast to previous reports(23), BRCA1 remained readily detectable on unsynapsed X and Y chromosome axes in mid-pachytene spermatocytes from *Atr^f/-^ Stra8-GFPCre* males, regardless of whether X-Y synapsis at the PAR was observed (Figure 6I-J). BRCA1 signals were also detected at the PAR in more than half of mid-pachytene spermatocytes (Figure 6I-J), consistent with previous observations(23). Therefore, BRCA1 localization to unsynapsed chromosome axes is independent of ATR in both zygotene and mid-pachytene spermatocytes, arguing against reciprocal regulation and further supporting a BRCA1-ATR regulatory hierarchy at unsynapsed chromosome axes during male meiotic prophase.

## Discussion

Our study identifies ATR as a master regulator of the oocyte meiotic prophase checkpoint. This conclusion is most strongly supported by the complete rescue of the survival of *Dmc1^-/-^* oocytes, which exhibit severe meiotic recombination defects characterized by persistent DSBs and extensive chromosome asynapsis. Despite these abnormalities, ATR loss essentially abolishes oocyte elimination, indicating that ATR is required for nearly the entire checkpoint response in *Dmc1^-/-^* oocytes. Notably, *Atr^-/-^* rescues *Dmc1^-/-^* oocytes more effectively than *Brca1^-/-^*, further supporting a central role for ATR in the oocyte meiotic prophase checkpoint.

The essential role of ATR in the chromosome asynapsis checkpoint is more directly demonstrated by the full rescue of the survival of *Spo11^-/-^* oocytes and further supported by the full rescue of the survival of *Trip13^-/-^* oocytes. Together with our previous studies(13,18), these findings establish a BRCA1-ATR signaling axis that links chromosome asynapsis to oocyte elimination. Importantly, BRCA1 is required for ATR localization to unsynapsed chromosome axes in both oocytes and spermatocytes, whereas ATR is dispensable for BRCA1 localization, arguing against the previously proposed reciprocal regulation and suggesting a conserved mode of BRCA1-ATR regulation in both sexes(23). Nevertheless, the more severe defect in recombinase loading caused by BRCA1 loss than by ATR loss suggests that BRCA1 also has ATR-independent functions during meiotic recombination. How BRCA1 regulates ATR localization to unsynapsed chromosome axes remains unclear. Interestingly, *Atr^-/-^* rescues *Spo11^-/-^*and *Trip13^-/-^* oocytes more effectively than *Brca1^-/-^*, raising the possibility that BRCA1 is not the sole factor regulating ATR localization to unsynapsed chromosome axes. A BRCA1-independent mechanism may therefore mediate residual ATR localization in the absence of BRCA1, with residual ATR activity sufficient to trigger elimination in a subset of oocytes.

An important implication of the *Trip13^-/-^* model is that the chromosome asynapsis checkpoint must be properly terminated once homologous chromosomes are fully synapsed. We previously showed that DSB repair is completed before *Trip13^-/-^* oocytes are eliminated(18), indicating that persistent DSBs do not account for their elimination. Instead, HORMAD1, HORMAD2, and BRCA1 remain aberrantly retained on fully synapsed chromosome axes. Here, we show that ATR is also aberrantly retained on synapsed chromosome axes in a BRCA1-dependent manner. Together, these findings indicate that persistent chromosome asynapsis checkpoint signaling after chromosome synapsis can drive inappropriate oocyte elimination. The strong dependence of the γH2AX signal on ATR in *Trip13^-/-^* oocytes further indicates that γH2AX in this context primarily reflects aberrant checkpoint signaling rather than persistent DSBs. Therefore, γH2AX should not necessarily be interpreted as evidence of unrepaired DNA damage during meiotic prophase. These findings highlight the importance of timely removal of checkpoint proteins from synapsed chromosome axes in terminating the chromosome asynapsis checkpoint.

An unresolved question is whether persistent DSBs and chromosome asynapsis activate independent or overlapping checkpoint responses during oocyte meiotic prophase. Despite its essential role in the oocyte meiotic prophase checkpoint, ATR is dispensable for oocyte elimination in response to IR-induced DNA damage. We previously found that BRCA1 is similarly dispensable for IR-induced oocyte elimination(13). However, one important distinction is that SPO11-induced programmed DSBs are generated and processed on chromosome axes, whereas IR-induced DNA damage is not restricted to chromosome axes. Therefore, the dispensability of ATR and BRCA1 for IR-induced oocyte elimination does not exclude the possibility that they participate in checkpoint signaling triggered by persistent DSBs. Although the *Dmc1^-/-^* model cannot distinguish the effects of persistent DSBs from chromosome asynapsis, the complete rescue by *Atr^-/-^* indicates that, if persistent DSBs contribute to the elimination of *Dmc1^-/-^* oocytes, this checkpoint response also requires ATR. Although ATR is not specifically localized to DSBs, it localizes along chromosome axes that contain DSB sites, providing a potential basis for signaling from persistent DSBs to ATR. Such signaling could share components with the chromosome asynapsis checkpoint, including BRCA1, or could involve distinct pathways that converge on ATR. Whether checkpoint signaling arising from persistent meiotic DSBs uses mechanisms distinct from or overlapping with those of the chromosome asynapsis checkpoint remains to be determined.

## Materials and Methods

### Mice

*Atr^f/f^* mice were obtained from The Jackson Laboratory (Strain No. 030135). *Chk2^−/−^* mice were obtained from GemPharmatech (Strain No. T027449). *Stra8-GFPCre* mice were a kind gift from Ming-Han Tong (Chinese Academy of Sciences)(30). *Dmc1^−/−^* mice were a kind gift from Hongbin Liu (Shandong University)(31). *Spo11^−/−^* mice were a kind gift from Qinghua Shi (University of Science and Technology of China)(32). *Brca1^f/-^ Stra8-GFPCre*, *Trip13^−/−^*, and *Brca1^−/−^ 53bp1^−/−^* mice were described previously(13,18,29).

Mice were genotyped using DNA extracted from tails or toes. Primer sequences for mouse genotyping are shown in Table S1. All experimental procedures were approved by Zhejiang University Animal Care and Use Committee (ZJU20190144, ZJU20220228 and ZJU20250615). All mice were housed under controlled environmental conditions (temperature 20-22°C, humidity 50-70% and 12 h light/12 h dark cycle) with *ad libitum* access to food and water.

### Preparation of meiotic spreads

For spreads of oocytes at the meiotic prophase, ovaries were isolated from fetal or newborn female mice. Ovaries were incubated in hypotonic extraction buffer (30 mM Tris-HCl pH 8.0, 50 mM sucrose, and 17 mM sodium citrate) for 25 min and then transferred into droplets of 100 mM sucrose buffer. Oocytes were released by chopping the ovaries in 100 mM sucrose buffer for 3 min. After removing large tissue fragments, the cell suspension was mixed with an equal volume of fixative buffer (1% PFA, 0.15% Triton X-100, and 10 mM borate buffer, pH 9.2). The mixture was smeared onto coverslips and air-dried.

For spreads of spermatocytes at meiotic prophase, testes were dissected from PD28 males. After removing the tunica albuginea, seminiferous tubules were digested in collagenase IV solution (1 mg/mL) at 37°C for 20 min. Cell pellets were resuspended in hypotonic extraction buffer (30 mM Tris-HCl pH 8.2, 50 mM sucrose, and 17 mM sodium citrate) for 10 min and then in 100 mM sucrose buffer for 5 min. The cell suspension was then mixed with an equal volume of fixative buffer, smeared onto coverslips and air-dried.

### Immunofluorescent (IF) staining and imaging

For IF staining of meiotic spreads, coverslips were treated with 0.5% Triton X-100 for 10 min and then incubated with primary antibodies at room temperature for 1 h. After washing in phosphate-buffered saline (PBS) three times, coverslips were incubated with secondary antibodies at room temperature for 1 h. Finally, coverslips were labeled with Hoechst 33342 solution for 5 min and then mounted with an anti-fade reagent. Images were captured using a fluorescence microscope with a CCD camera (ECLIPSE Ti2-E, Nikon).

For IF staining of oocytes, germinal-vesicle stage oocytes from PD21 female mice were harvested in M2 medium 44 h after PMSG injection and cultured in microdrops of M16 medium covered with mineral oil for an additional 8 h at 37 °C in a 5% CO2 atmosphere. IF staining of MI-stage oocytes was performed as described above. Images were taken using a laser scanning confocal microscope (STELLARIS 5, Leica Microsystems).

For IF staining of paraffin sections of ovaries, sections were deparaffinized, rehydrated and incubated with antigen retrieval buffer (10 mM sodium citrate with 0.05% Tween, pH 6.0) at 95 °C for 30 min. IF staining of ovarian sections was performed similarly as described above. Images were taken using a laser scanning confocal microscope (STELLARIS 5, Leica Microsystems). Antibody information is listed in Table S2.

### Histological analysis and follicle count

Ovaries, testes and epididymides were dissected and fixed in 4% PFA at 4°C overnight. Fixed tissues were dehydrated using increasing concentrations of ethanol and then embedded in paraffin for serial sectioning at a thickness of 5 μm. The sections were stained with hematoxylin and eosin (H&E). Follicle numbers in PD21 and PD60 ovaries were quantified by counting every fifth section. For follicle quantification of PD9 ovaries, MVH-positive cell numbers were counted in the central ovarian section. Only one ovary per animal was used for follicle quantification. Histological images were acquired using the Grundium Ocus microscope slide scanners (OCUS01, Grundium).

### Fertility test

*Atr^f/-^ Stra8-GFPCre* female mice and their corresponding control mice were mated with WT adult male mice once they reached 8 weeks of age. Litter sizes were determined by counting pups at birth for 3 consecutive months.

### Analysis of early embryonic development

PD21 female mice received intraperitoneal injections of 5 IU PMSG, followed 44 h later by 5 IU hCG. Immediately after the injection of hCG, female mice were placed individually with WT adult male mice. Successful mating was confirmed by the presence of vaginal plugs. For early embryonic development analysis, embryos were collected at 1.5 and 3.5 days after fertilization. Unfertilized oocytes and embryos that failed to reach the zygote stage were excluded from the analysis.

### Ionizing radiation (IR)

To investigate the IR sensitivity of oocytes, PD2 female mice were exposed to a single dose of 0.45 Gy IR and then their ovaries were isolated at PD9 for histological analysis.

### Statistical analysis

At least three independent samples were included in each experiment. Statistical analysis was performed using GraphPad Prism 9, and the results were presented as means ± SD. To calculate statistical significance between different groups, two-tailed unpaired Student’s t tests were performed.

## Acknowledgements

We thank Hongbin Liu, Qinghua Shi, and Ming-Han Tong for sharing mouse strains. We thank Quanfeng Zhu for helping with mouse genotyping. We thank Chao Bi and Xiaoli Hong from the Core Facilities, Zhejiang University School of Medicine for their technical support. We thank Xiaofen Jin, Yumiao Niu, and Feiting Xie from the Central Laboratory, Women’s Hospital, Zhejiang University School of Medicine for their technical support. This work is funded by National Natural Science Foundation of China (32370900 and 32570998 to L.L.).

## Author Contributions

Jiwei Li: Investigation, Formal analysis, Writing – original draft, Writing – review and editing. Xiaofei Jiao: Investigation, Formal analysis, Writing – review and editing. Long Bai: Investigation, Formal analysis, Writing – review and editing. Yuan Wang: Investigation. Yidan Liu: Conceptualization, Supervision, Writing – review and editing. Yimin Zhu: Conceptualization, Supervision, Writing – review and editing. Lin-Yu Lu: Conceptualization, Supervision, Writing – review and editing.

## Data Availability Statement

Data supporting this study are included within the article.

---

**Figure S1.**
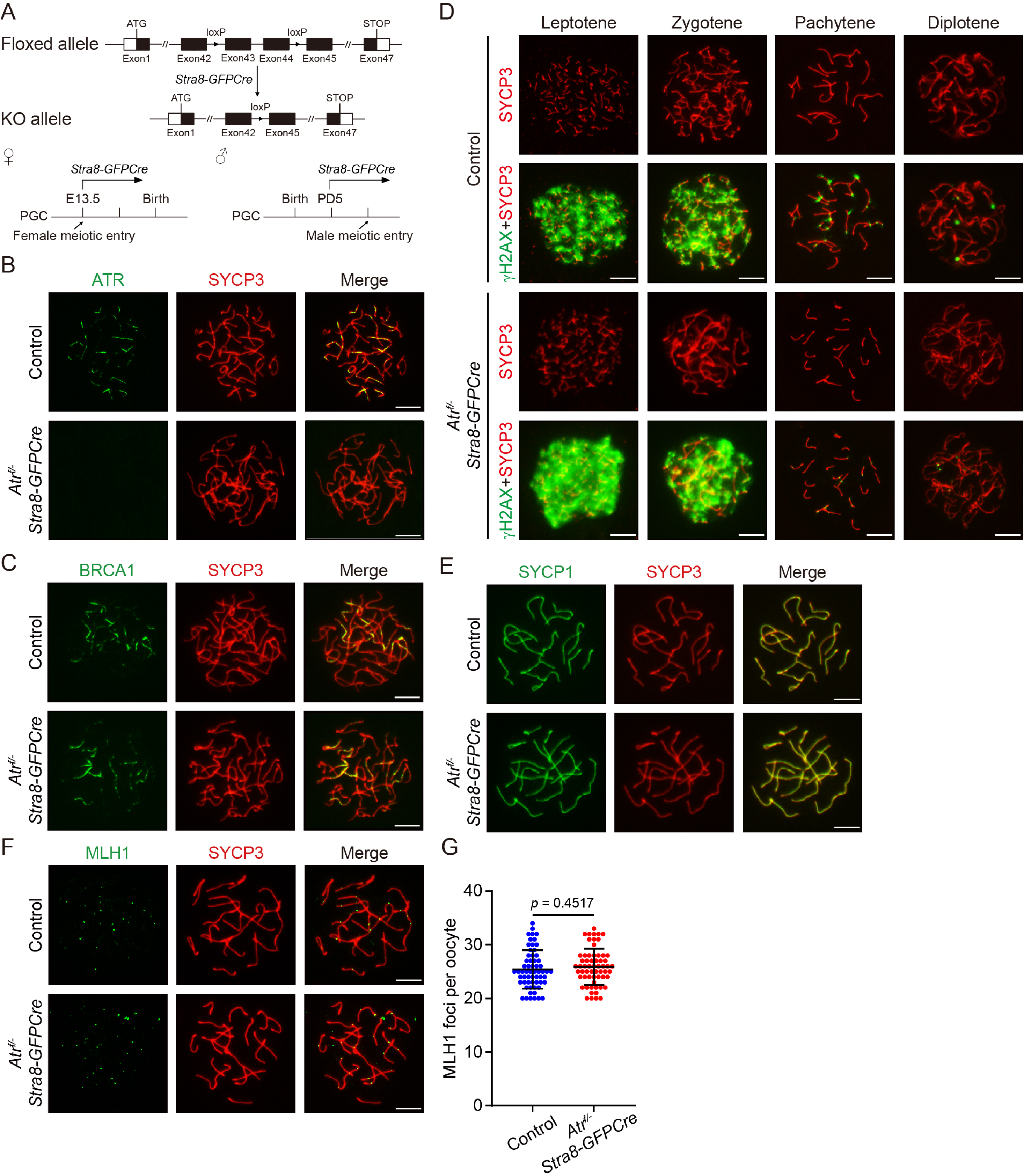
*Atr^f/-^ Stra8-GFPCre* oocytes exhibit largely normal meiotic recombination. (A) Schematic illustration for generation of *Atr^f/-^ Stra8-GFPCre* mice. (B, C) Representative images of ATR and SYCP3 staining (B) and BRCA1 and SYCP3 staining (C) in chromosome spreads of control and *Atr^f/-^ Stra8-GFPCre* oocytes at zygotene stages. Scale bars, 10 μm. (D) Representative images of γH2AX and SYCP3 staining in chromosome spreads of control and *Atr^f/-^ Stra8-GFPCre* oocytes at leptotene, zygotene, pachytene and diplotene stages. Scale bars, 10 μm. (E, F) Representative images of SYCP1 and SYCP3 staining (E) and MLH1 and SYCP3 staining (F) in chromosome spreads of control and *Atr^f/-^ Stra8-GFPCre* oocytes at pachytene stages. Scale bars, 10 μm. (G) Quantification of MLH1 foci in chromosome spreads of control and *Atr^f/-^ Stra8-GFPCre* oocytes. All statistical data are presented as means ± SD. *P* values were determined by two-tailed unpaired Student’s t tests.

**Figure S2.**
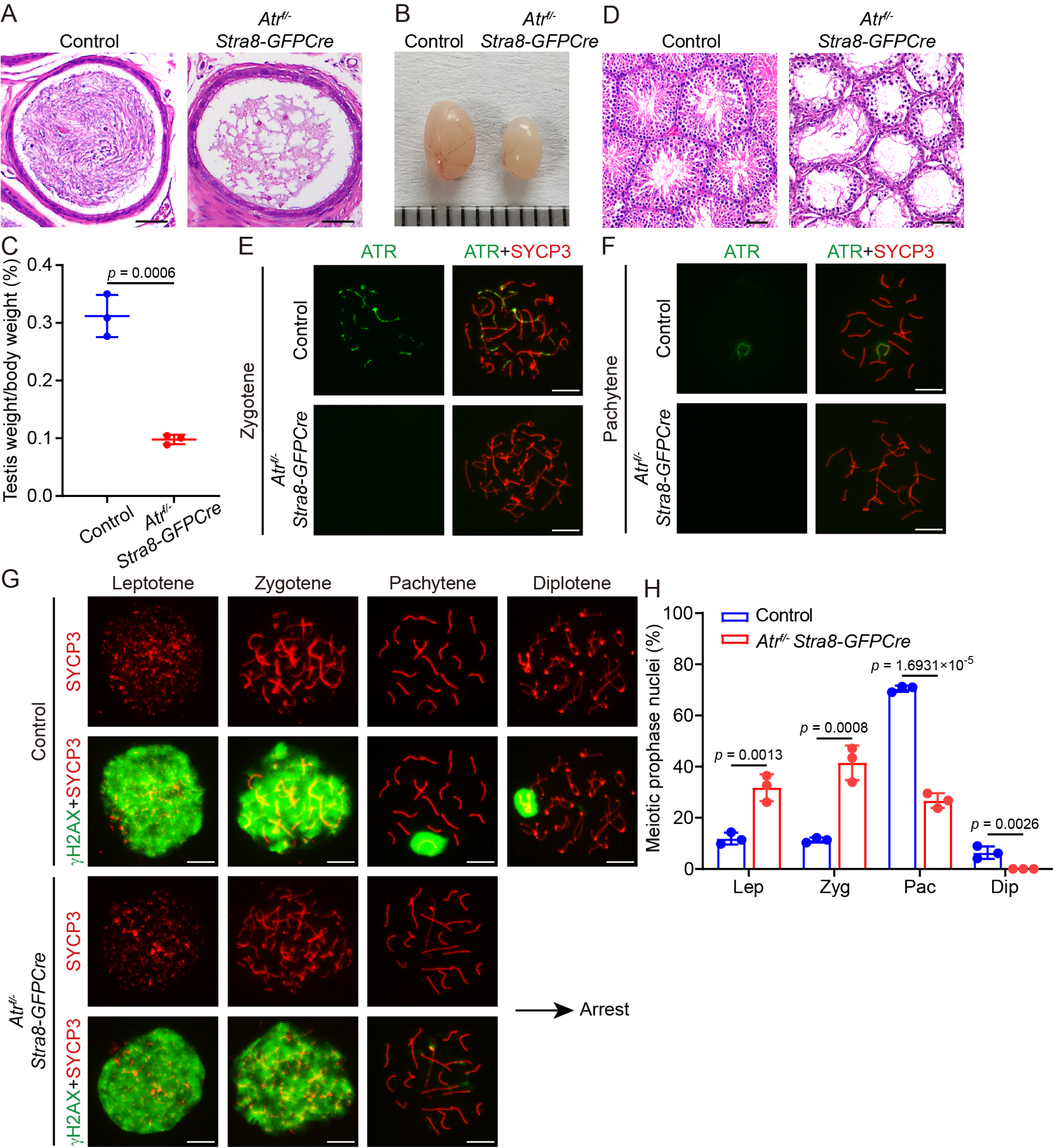
ATR is essential for male fertility. (A) Representative images of H&E staining of paraffin sections from control and *Atr^f/-^ Stra8-GFPCre* cauda epididymides at PD60. Scale bars, 100 μm. (B) Representative images of control and *Atr^f/-^ Stra8-GFPCre* testes at PD28. (C) The ratios of testis weight to body weight in control and *Atr^f/-^ Stra8-GFPCre* mice at PD28. (D) Representative images of H&E staining of paraffin sections from control and *Atr^f/-^ Stra8-GFPCre* testes at PD28. Scale bars, 100 μm. (E, F) Representative images of ATR and SYCP3 staining in chromosome spreads of control and *Atr^f/-^ Stra8-GFPCre* spermatocytes at zygotene (E) and pachytene (F) stages. Scale bars, 10 μm. (G) Representative images of γH2AX and SYCP3 staining in chromosome spreads of control and *Atr^f/-^ Stra8- GFPCre* spermatocytes at leptotene, zygotene, pachytene, and diplotene stages. Scale bars, 10 μm. (H) Ratios of control and *Atr^f/-^ Stra8-GFPCre* spermatocytes at each stage of meiotic prophase. All statistical data are presented as means ± SD. *P* values were determined by two-tailed unpaired Student’s t tests.

**Figure S3.**
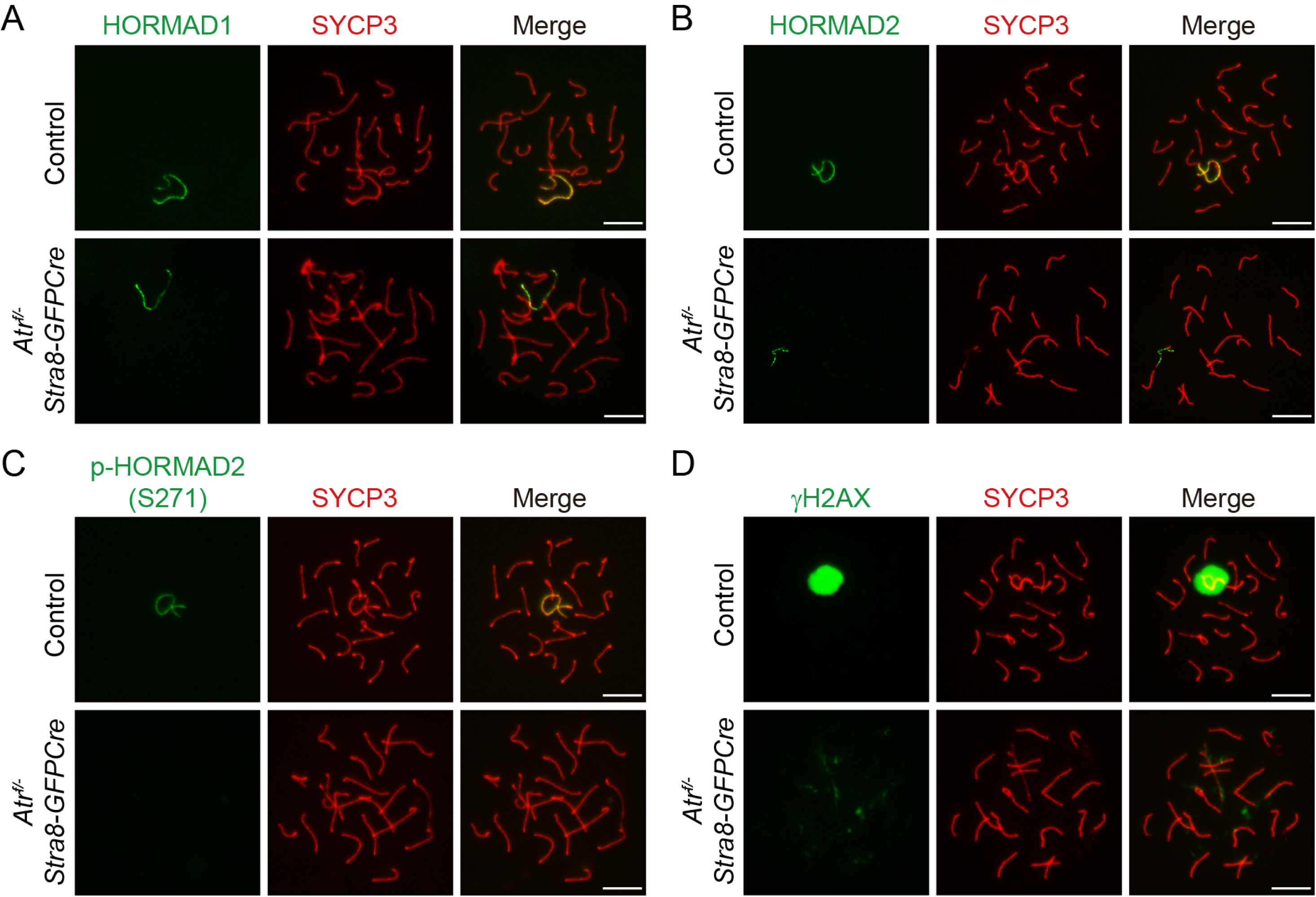
ATR regulates meiotic sex chromosome signaling in spermatocytes. (A-D) Representative images of HORMAD1 and SYCP3 staining (A), HORMAD2 and SYCP3 staining (B), p-HORMAD2 (S271) and SYCP3 staining (C) and γH2AX and SYCP3 staining (D) in chromosome spreads of control and *Atr^f/-^ Stra8-GFPCre* spermatocytes at pachytene stages. Scale bars, 10 μm.

**Table S1.**
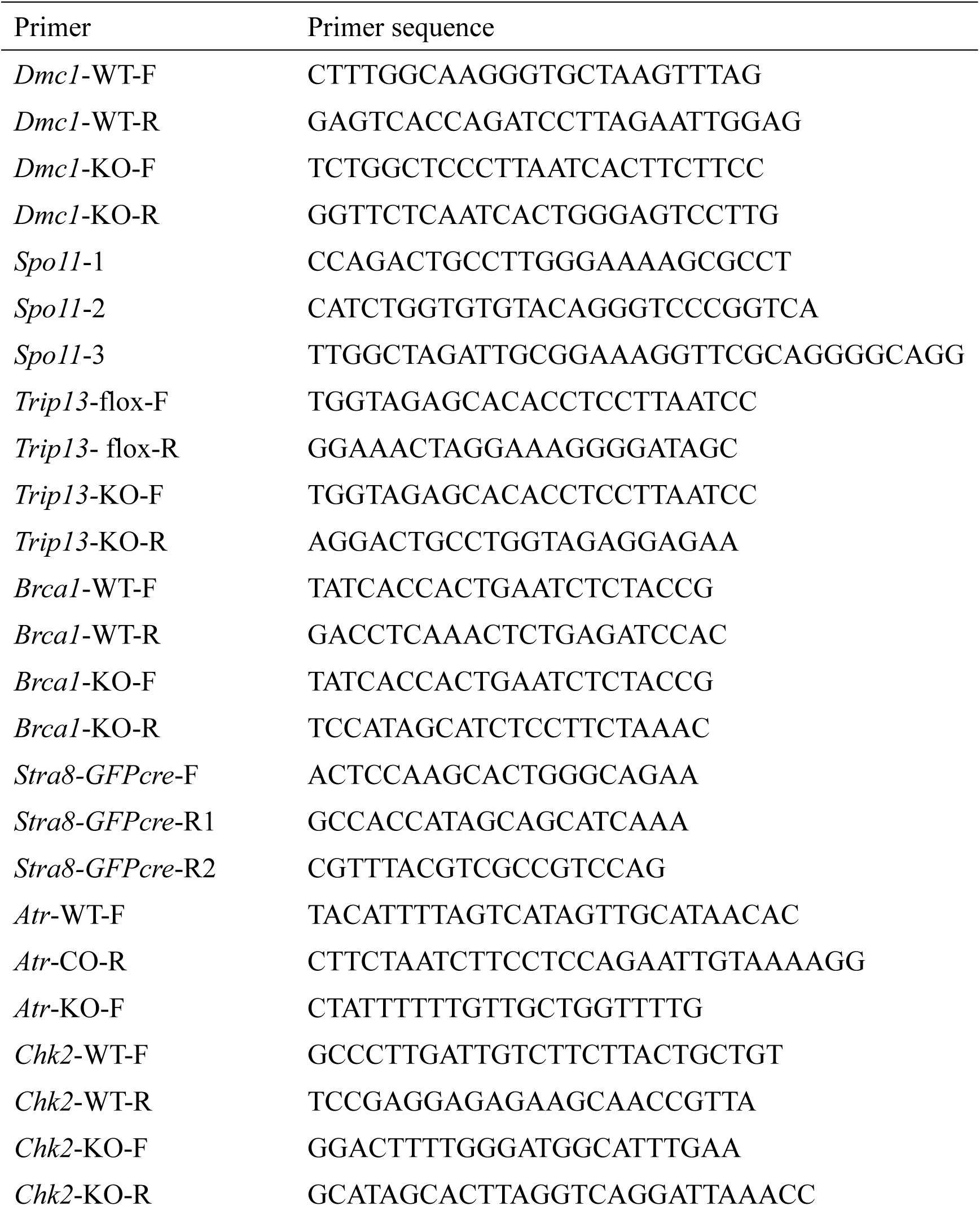
Primers used for genotyping in this study.

**Table S2.**
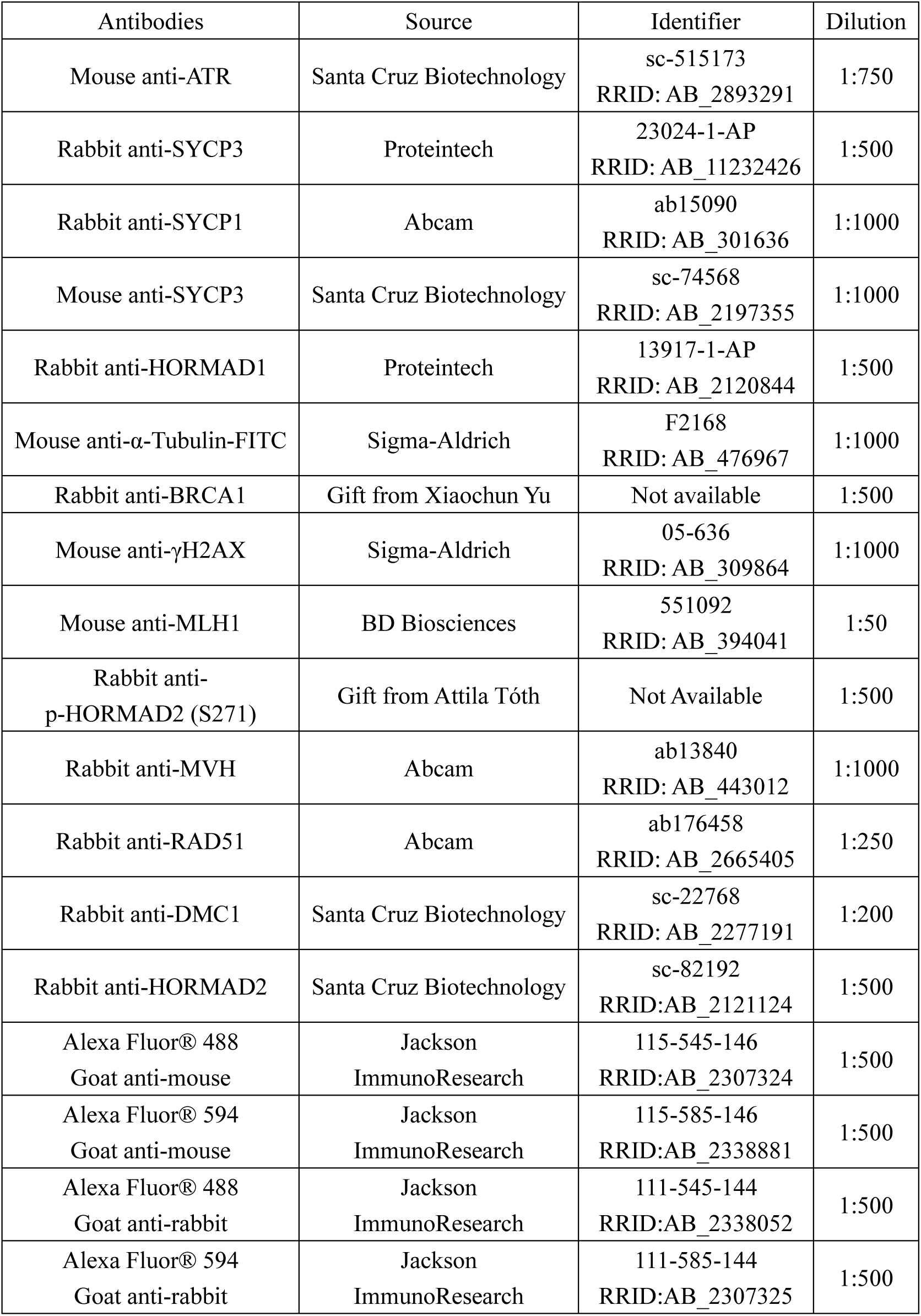
Antibodies used for immunofluorescence staining in this study.

